# Royalactin induces copious longevity via increased translation and proteasome activity in *C*. *elegans*

**DOI:** 10.1101/421818

**Authors:** Giel Detienne, Pieter Van de Walle, Wouter De Haes, Bram Cockx, Bart P. Braeckman, Liliane Schoofs, Liesbet Temmerman

## Abstract

As demonstrated in various animal models, organismal longevity can be achieved via interventions that at the mechanistic level could be considered to entail ‘defensive’ responses: most long-lived mutants focus on somatic maintenance, while reducing growth pathway signalling and protein translation and turnover. We here provide evidence that the opposite mechanism can also lead to longevity and improved health.

We report on the mode of action of royalactin, a glycoprotein activator of epidermal growth factor signalling, capable of extending lifespan in several animals. We show that in *Caenorhabditis elegans*, royalactin-induced longevity depends on increased protein translation and entails increased proteasome activity. We propose the term ‘copious longevity’ to describe this newly-elucidated mechanism. In contrast to what is true for many other lifespan-extending interventions, we observed no obvious trade-offs between royalactin-induced longevity and several life history traits. Our data point towards increased protein turnover to support healthy ageing, and provide a means for future comparative studies of defensive vs. copious mechanisms.

## Introduction

When studying the trajectories along which organisms age, one may wonder whether longevity can be achieved free from trade-offs at all. For example, growth and to a lesser extent reproduction are often inversely correlated with lifespan^1,2^. However, some species escape this general rule. One popular example are eusocial animals – such as the honeybee *Apis mellifera* – in which the highly reproductively active queen caste grow larger and can live up to ten times longer than the sterile worker caste^3^. These organisms may therefore unveil mechanisms underlying an ageing trajectory that escapes some of the general trade-offs associated with longevity.

We here address this broader question by studying the mode of action of the bee-specific pro-longevity compound royalactin. In *A. mellifera*, royalactin stimulates the epidermal growth factor (EGF) signalling pathway and ultimately leads to epigenetic changes and a long-lived queen phenotype^4^. Ingestion of royalactin induces proliferative effects and health benefits that appear to be conserved across species^4,5^. We previously demonstrated that also in the popular ageing model *Caenorhabditis elegans*, royalactin requires EGF and its receptor for inducing longevity, providing the first evidence that royalactin can prolong lifespan outside the order of insects^6^. Royalactin also enhances exogenous oxidative stress tolerance^7^ and locomotion^6^ in adult *C. elegans*, suggesting a positive effect on healthspan as well. Yet, the mechanism by which these pro-longevity effects are achieved, is still elusive.

Our data point towards an unexpected mechanism that supports healthy ageing via rebuilding of cellular components, rather than the more commonly observed mechanism^8–10^ of stabilizing the existing proteome.

## Results

### Royalactin-mediated lifespan extension is incompatible with decreased protein translation

To investigate how royalactin - and indirectly so EGF receptor (EGFR) signalling - is able to extend lifespan, we performed a differential proteomics experiment in *C. elegans* (see also Supplementary Information). The various proteins that are upregulated after royalactin treatment of wild-type (WT) *C. elegans* are associated with protein translation, ribosomal function, cellular detoxification and muscle maintenance, or possess a protein binding and folding (*e.g*. chaperone) function (Supplementary Dataset 1). Clustering of the differential proteins based on protein-protein interactions shows significant reconfiguration of the proteome towards increased protein translation, elongation and proteostasis (Supplementary Fig. 1). The implied increase in translation is counter-intuitive since many studies report lowered protein translation in long-lived animals^11–19^. Our data suggest a potentially different molecular mechanism for royalactin-mediated longevity; indeed, the bulk of protein abundance changes observed in royalactin-treated worms are opposed to those observed in most long-lived *C. elegans* (Supplementary Table 1). Most prominently, the cellular changes related to protein synthesis induced by royalactin (*e.g*. upregulation of elongation factors and chaperonins) are clearly opposite to those during dietary restriction, reduced insulin signalling and others, but predictably similar to those in long-lived EGFR gain-of-function mutants^20^ (Supplementary Table 1).

To test for dependency of royalactin-induced longevity on increased or dynamic translation, we assessed whether royalactin is still capable of extending *C. elegans* lifespan after reducing protein translation via genetic or chemical interventions. One of the main cellular regulators of protein translation is the p70 ribosomal S6 kinase – encoded by the *rsks*-*1* gene in *C. elegans* – which regulates protein translation via phosphorylation of the ribosomal protein S6^21^. *rsks*-*1* knockout reduces protein translation rates of adults by circa one third^12^ and as expected^12,15,22^, also clearly extended lifespan by over a third compared to WT (Fig. 1A, Supplementary Dataset 2). Whereas WT animals displayed enhanced survival after royalactin treatment, *rsks*-*1(*-) animals show a significant reduction in survival when fed royalactin (Fig. 1A, Supplementary Dataset 2).

**Fig. 1:**
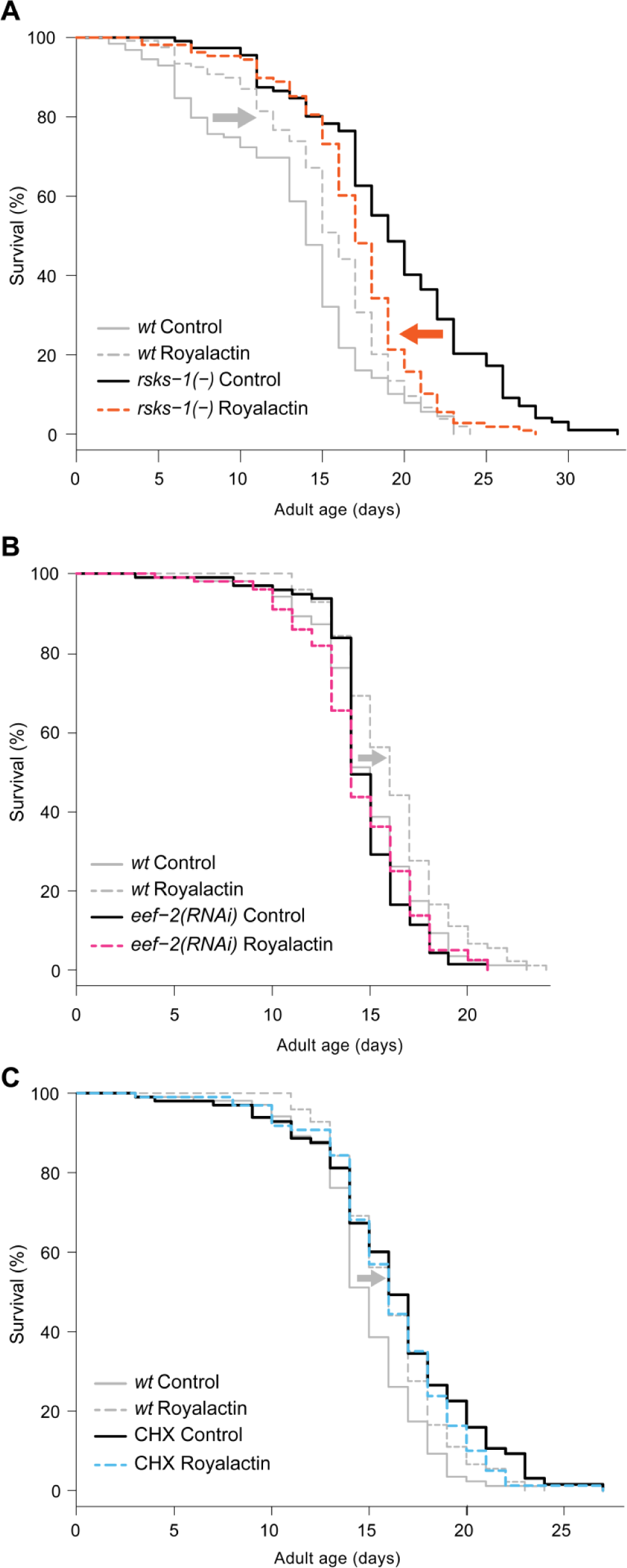
Reduction of protein translation via genetic or chemical interventions abolishes the lifespan-extending effect of royalactin in *C. elegans.* Survival curves under three conditions of reduced protein translation, in the presence and absence of royalactin. We observed a clearly diminished effect of royalactin on survival after reducing protein translation, compared to the effect of royalactin in control WT animals. **(A)** Genetic knockout of the p70 S6 kinase RSKS-1. The grey and orange arrows depict the change in survival after royalactin treatment in the WT control condition and during RSKS-1 knockout, respectively. **(B)** Knockdown of eukaryotic elongation factor EEF-2 via RNAi in adults. **(C)** Protein translation restriction via the addition of 2 μM cycloheximide (CHX) from the late L4 stage (Detailed lifespan data: Supplementary Dataset 2).

As a second genetic intervention to reduce mRNA translation, we knocked down the eukaryotic translation elongation factor 2 (*eef*-*2*). In our hands, this did not result in a significant decline in survival compared to control animals (Fig. 1B, Supplementary Dataset 2). The survival decrease we observed was less pronounced than a previous observation by Dong and colleagues^23^, presumably due to differences in the exact timing of the experiment and/or media composition^24,25^. When *eef*-*2* was knocked down, royalactin was no longer capable of extending lifespan of *C. elegans* (Fig. 1B, Supplementary Dataset 2), in contrast to royalactin-fed controls that showed significantly increased survival (Fig. 1B, Supplementary Dataset 2).

Next, we directly blocked translation elongation by administering cycloheximide (CHX) to *C. elegans.* The relatively low CHX concentration we used (2 μM) is known to block the translational elongation step in cytoplasmic, but not mitochondrial protein synthesis by binding to the 60S ribosomal subunit and inhibiting EEF-2-guided translocation^26,27^. Such CHX treatment has been reported to extend lifespan of *C. elegans* before^28^, and largely mimics the effect of the direct knockdown of translation elongation factors^24^. Addition of CHX to the growth medium from the late L4 stage onwards resulted in a significant increase in survival (Fig. 1C, Supplementary Dataset 1) compared to survival on control medium. In contrast to WT controls, royalactin could not extend the lifespan of CHX-treated animals (Fig. 1C, Supplementary Dataset 1).

Taken together, our data imply the necessity of protein translation in royalactin-induced longevity, supporting the notion that royalactin-mediated longevity relies on enhanced or at least dynamic protein translation (Fig. 1).

### Royalactin increases translation and promotes proteasome function

Our survival (Fig. 1) and proteomics (Supplementary Fig. 1) data point towards dynamic translation as being an integral part of royalactin-mediated longevity. To further substantiate that mRNA translation is increased, we performed polysome profiling. This method quantifies the amount of actively translated mRNA molecules in an extract^12,29^. We observed an increase in monosome and polysome peak areas for royalactin-treated animals (Fig. 2, Supplementary Dataset 3) of respectively 26.2% and 29.8%. Therefore, per unit of mRNA present in the cytosolic lysates, mRNA from the royalactin-treated group is more likely to be translationally active^30^. The polysome/monosome ratio remained stable between both groups (Fig. 3) and we did not detect any significant changes in total amount of mRNA present in the initial lysates (Supplementary Dataset 3).

**Fig. 2:**
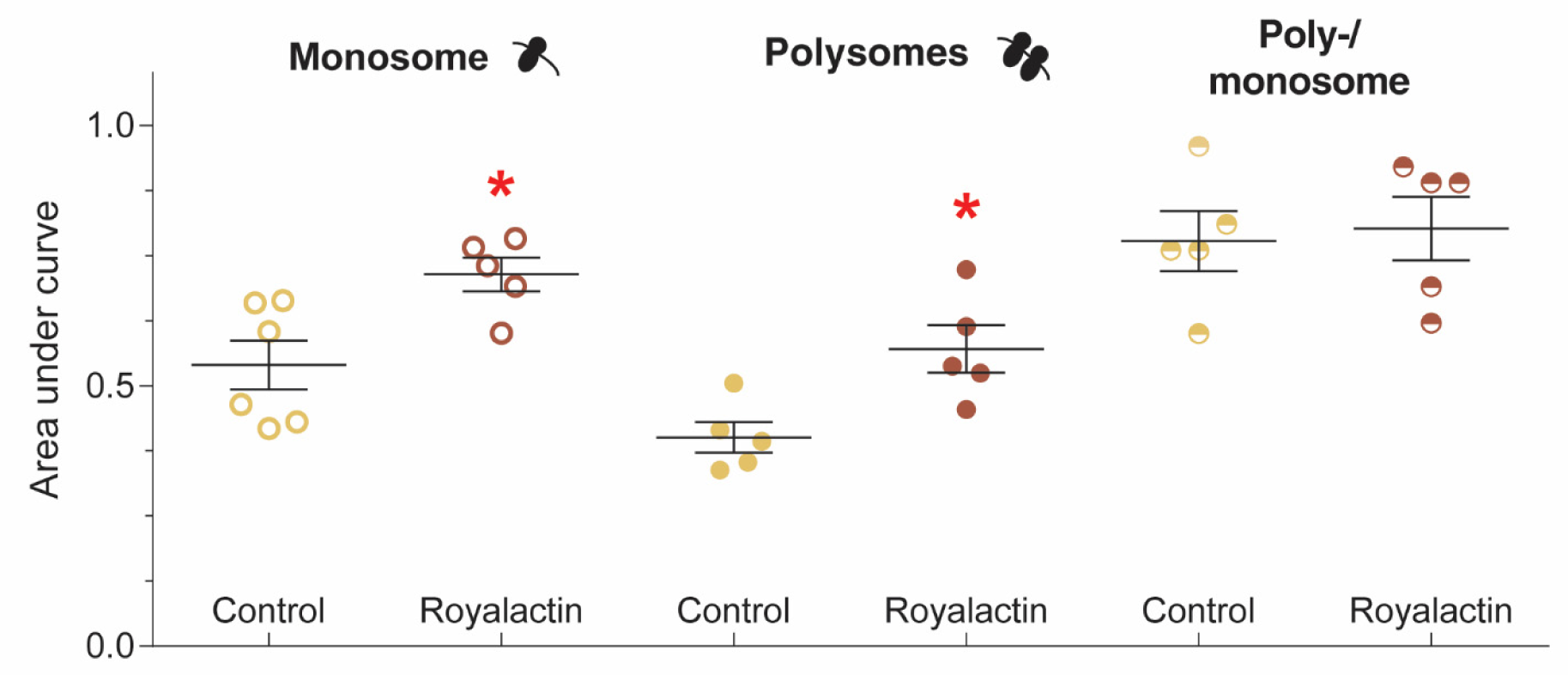
Polysome profiling indicates an overall proportional increase in mRNA translation in royalactin-treated animals. The monosome and polysome fractions are more abundant in royalactin-treated animals, while the polysome/monosome ratio remains unchanged. See Supplementary Dataset 3 for more detailed results and representative absorbance profiles. Error bars indicate SEM; ^∗^ *p*_*t*-*test*, *FDR*_ < 0.05. Averages ± SEM from left to right: 0.54 ± 0.047, 0.71 ± 0.072, 0.40 ± 0.029, 0.57 ± 0.046, 0.78 ± 0.058, 0.80 ± 0.061.

**Fig. 3:**
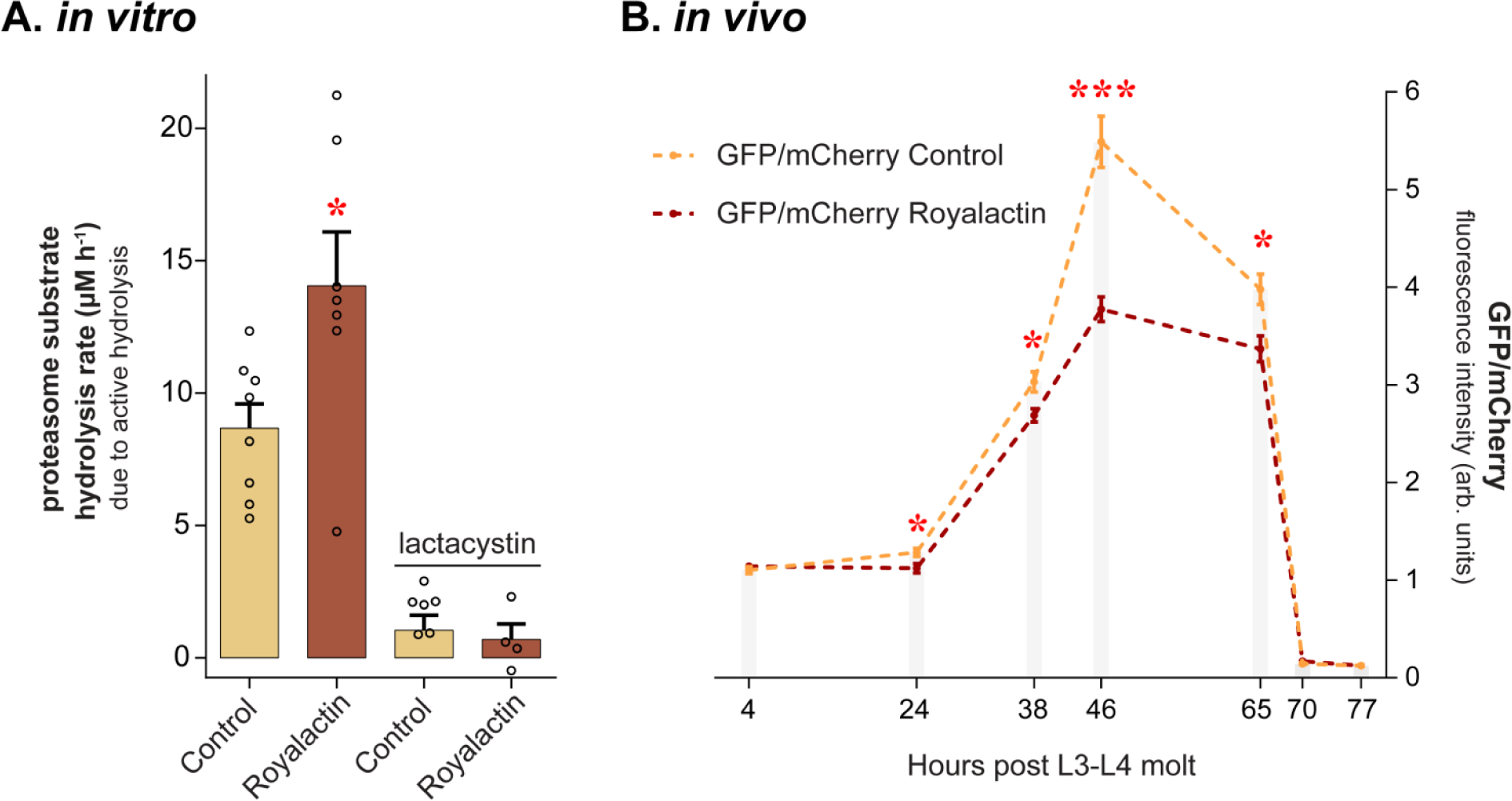
Royalactin enhances proteasome activity *in vitro* and *in vivo.* **(A)** Organism-wide 20S proteasome activity *in vitro* is enhanced after royalactin treatment of *C. elegans* (*p*_*log*-*rank*_ = 0.025). The specific proteasome inhibitor lactacystin abolishes the increase in chymotryptic substrate hydrolysis rate in royalactin-fed animals. Individual biological replicates, each derived from thousands of animals, are indicated with hollow dots. Error bars indicate SEM; ^∗^ *p*_*log*-*rank*_ < 0.05. Averages ± SEM from left to right: 8.67 ± 0.92; 14.06 ± 2.02; 1.05 ± 0.56; 0.70 ± 0.59. **(B)** The turnover of GFP marked for UPS-based degradation is enhanced in royalactin-treated *C. elegans*, indicating enhanced *in vivo* UPS activity. Fluorescence from non-modified mCherry driven from the same promoter was used to correct for differences in translation. Error bars indicate SEM; ^∗^ *p*_*t*-*test*,*FDR*_ < 0.05; ^∗∗∗^ *p*_*t*-*test*,*FDR*_ < 0.001. Averages ± SEM and the amount of biological replicates from left to right, starting with the control condition: 1.12 ± 0.037 (12), 1.14 ± 0.027 (11), 1.29 ± 0.044 (11), 1.12 ± 0.048 (11), 3.03 ± 0.11 (11), 2.69 ± 0.07 (11), 5.49 ± 0.26 (17), 3.72 ± 0.14 (15), 4.06 ± 0.19 (14); 3.37 ± 0.13 (22), 0.14 ± 0.0074 (27), 0.17 ± 0.015 (27), 0.13 ± 0.0061 (24), 0.14 ± 0.01 (26).

Since delaying the ageing-related collapse of proteostasis is considered to be a prerequisite for longevity^31^, it can now be asked whether the observed increase in translation is balanced by an increase in protein degradation. The major proteolytic system in the eukaryotic cytosol is the ubiquitin-proteasome system (UPS), which selectively degrades damaged, unfolded or misfolded proteins, as well as short-lived regulatory proteins^32^. We suspected UPS activation in *C. elegans* after royalactin treatment, since UPS activity generally correlates well with lifespan and stress resistance of organisms^20,32–35^ and has been linked to augmented EGF signalling before^20,36^. Indeed, *in vitro* 20S proteasome activity in homogenates of royalactin-fed animals revealed a significant increase in mean hydrolysis rate compared to that of homogenates of untreated animals (+62%, Fig. 3A). This increase was abolished when samples were mixed with lactacystin (Fig. 3A), a selective proteasome inhibitor leaving only non-proteasomal chymotrypsin activity (*e.g.* due to proteases in the gut or other tissues)^37^, indicating a specific increase in proteasome activity. We confirmed these results *in vivo*, where differential UPS activity of royalactin-treated animals peaked at 46 hours (*i.e.* at the young adult stage, 32% difference with the negative control), followed by an expected decline in fluorescence levels (Fig. 3B, Supplementary Fig. 3) due to a sustained GFP breakdown by the proteasome^38^.

Combined, our data support the notion that royalactin-mediated longevity relies on enhanced protein translation and degradation.

### The royalactin regimen is free of trade-offs for all tested traits

To assess functional impact of royalactin-induced longevity at the organismal level, we explored potential trade-offs with several life history and physiological parameters. These include reproductive fitness, body size and resilience against exogenous stressors. Each parameter is linked to hits in our proteomics screen, and/or to observations in other long-lived animals.

Total reproductive capacity or the temporal pattern thereof are often^39,40^ – though certainly not always [*e.g*.^41^] – altered in long-lived *C. elegans.* For royalactin, it would be reasonable to predict a positive effect on growth, reproduction and lifespan alike, since this is typically seen in long-lived castes of social insects^4^. None of the observed time points displayed a significant difference between progeny production of untreated and royalactin-treated animals (Fig. 4A, Supplementary Dataset 4), nor was there a significant difference between the total number of progeny (Fig. 4B). Therefore, under optimal conditions, there is no change in self-reproductive fitness of *C. elegans* due to royalactin treatment. Body size measurements of royalactin-treated and untreated worms led to similar results: average body length and body width increased, but only slightly, with respectively 3% and 2% (Supplementary Fig. 2). As such, we did not detect trade-offs between these life-history traits and royalactin-mediated longevity.

**Fig. 4:**
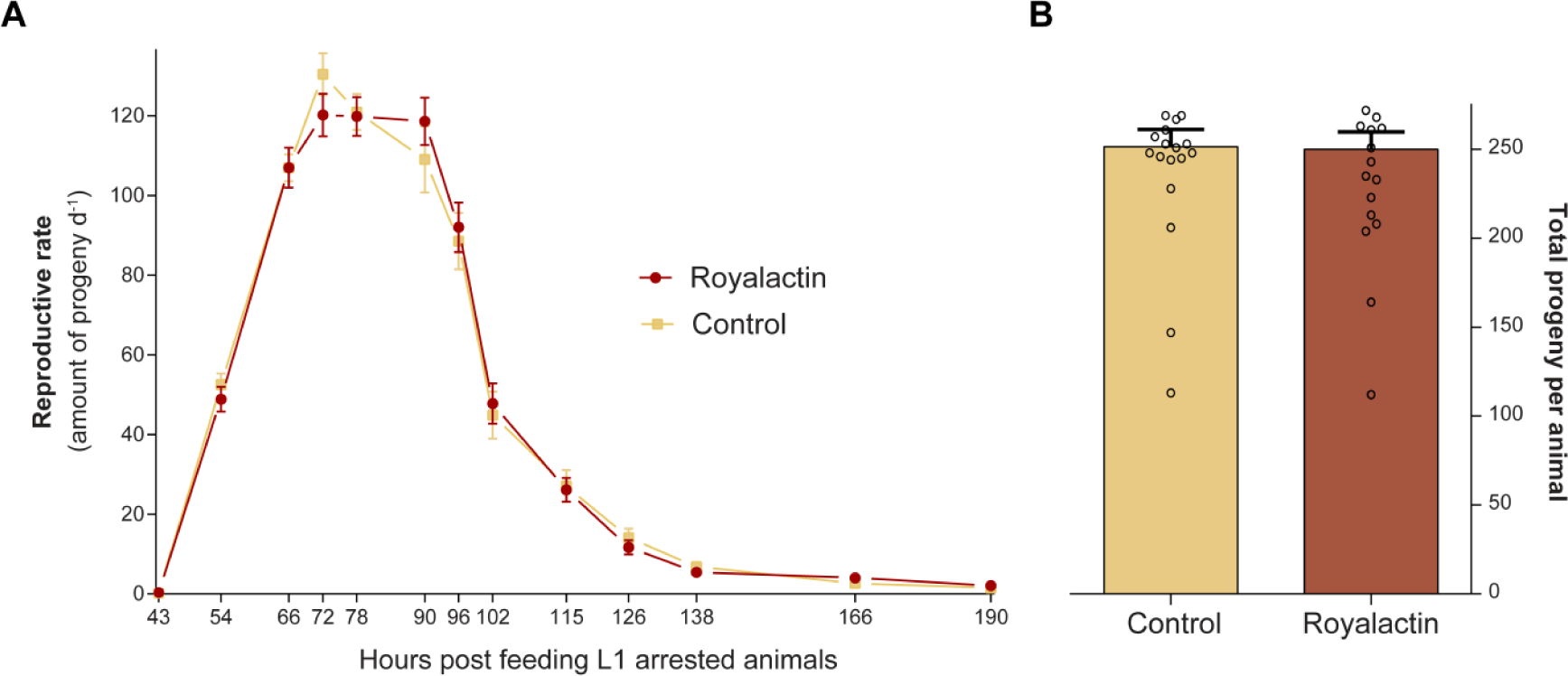
Royalactin treatment induces no significant change in self-reproductive fitness of WT *C. elegans.* Both reproductive span **(A)** and fecundity **(B)** are largely identical in WT royalactin-treated and untreated animals. Error bars indicate SEM. Individual biological replicates are indicated with hollow dots. Detailed data can be found in Supplementary Dataset 4.

We also monitored survival of *C. elegans* grown in the presence and absence of royalactin, under various acute stressors (Fig. 5, Supplementary Dataset 5), and measured transcription of stress-responsive GFP reporters^42^ to assess basal cellular stress levels after royalactin feeding. Our stress assays clearly showed that royalactin increases resistance to a variety of biotic and abiotic stressors (Fig. 5A-C, Supplementary Dataset 5). In line with our earlier observations for hydrogen peroxide resistance^7^, royalactin-mediated oxidative stress resistance to paraquat depends on the EGF-regulated EOR-1 transcription factor and its downstream target GST-4 (Fig. 5C). Combined with the observation that several fluorescent stress reporters were downregulated after royalactin treatment (Fig. 5D-F), we can conclude that generally, royalactin-treated worms are less stress-sensitive than controls (see also Supplementary information for a detailed exposition of stress assay results). To more specifically test enhanced resilience to proteostatic insults after royalactin treatment, we examined *C. elegans* expressing aggregation-prone polyglutamine (polyQ) repeats^43^. We observed a slight decrease in polyQ aggregates abundance from the L4 stage onwards in royalactin-treated animals (Fig. 5G), with the strongest effect around the late L4 stage (−22.3%).

**Fig. 5:**
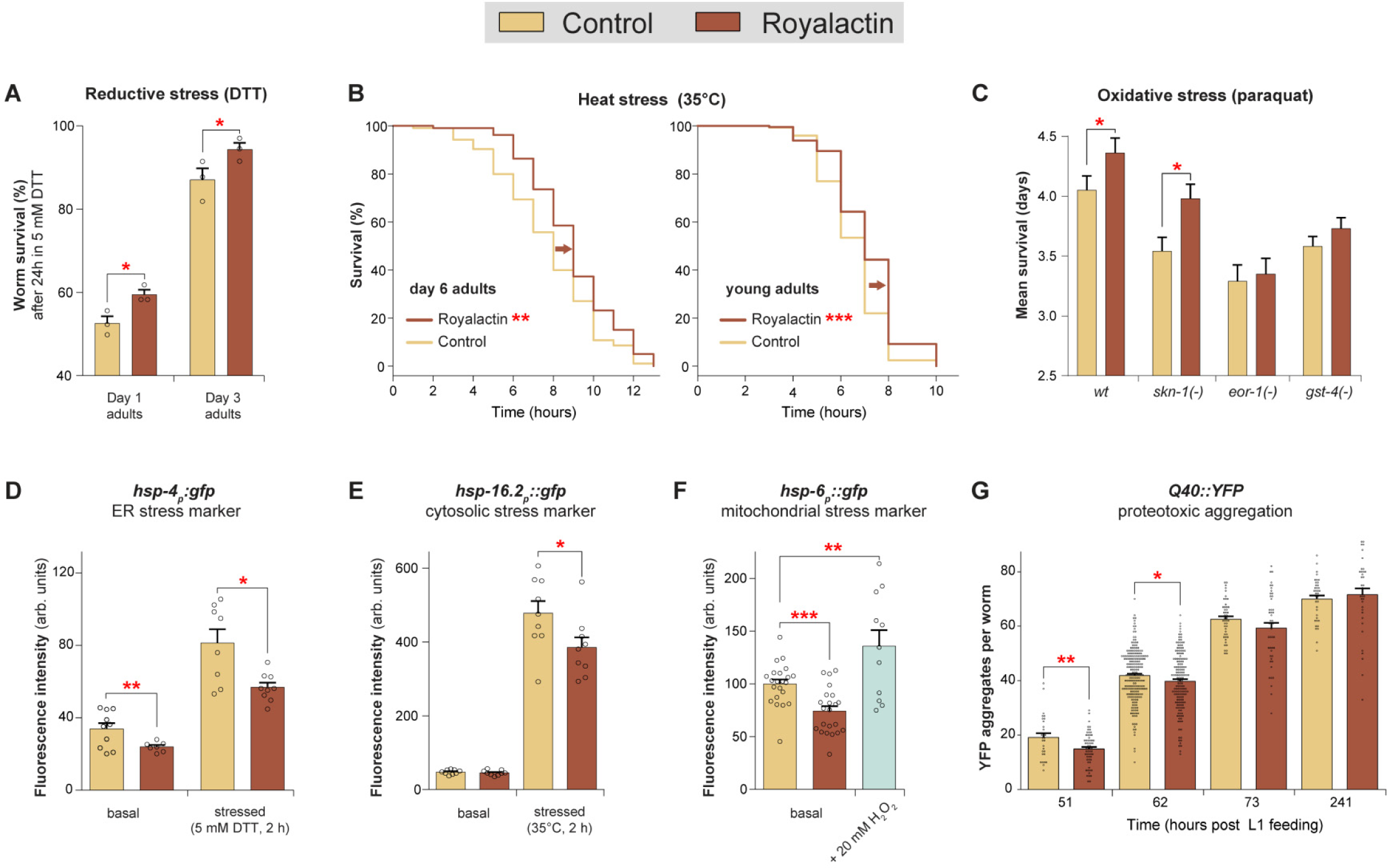
Royalactin-fed *C. elegans* survive several stressors better than untreated animals, display lower levels of stress-responsive GFP reporter transcription and a delay in polyQ aggregation. **(A)** Survival in 5 mM DTT is enhanced in royalactin-fed animals at days 1 and 3 of adulthood. ^∗^ *p*_*t*-*test*_ < 0.05 **(B)** Heat stress survival is significantly enhanced after royalactin treatment in day 6 WT adults (+15.1%, *p*_*log*-*rank*_ = 0.0035) and young adult worms (+13.8%, *p*_*log*-*rank*_ = 5.32E-9). **(C)** Survival on paraquat, a mitochondrial ROS inducer, is slightly but significantly increased in WT (+7.7%, *p*_*log*-*rank*_ = 0.048) and *skn*-*l(*-) animals (+12.4%, *p*_*log*-*rank*_ = 0.0117) treated with royalactin, while the effect diminishes and becomes insignificant when knocking out *eor-1* (+1.8% survival, *p*_*log*-*rank*_ = 0.718) or *gst*-*4* (+4.2%, *p*_*log*-*rank*_ = 0.215). (D) Mean *hsp*-*4p::gfp* fluorescence is significantly lower in royalactin-fed worms compared to controls, both under basal (−14.8%, *p*_*t*-*test*_ = 0.0034) and stressed conditions (−30.1%, *p*_*t*-*test*_ = 0.015). **(E)** Under stressed conditions, *hsp*-*l6.2p::gfp* fluorescence was significantly lower in royalactin-treated compared to control animals (−24.2%, *pANOVA*,*FDR* = 0.0086). Under basal conditions, fluorescence is low and similar for royalactin-treated and untreated worms (*p*_*ANOVA*,*FDR*_ = 0.94). **(F)** Expression of *hsp*-*6p::gfp*, a mitochondrial stress sensor, was decreased by 25.7% during royalactin-feeding compared to untreated animals (*p*_*t*-*test*_ = 0.0002). **(G)** Royalactin-fed worms displayed significantly lower levels of polyQ aggregates at 51 h (late L4 stage) and 62 h (young adult stage). Error bars always indicate SEM. See Supplementary Information for a more detailed description of results and Supplementary Dataset 5 for more information concerning the pooling of data, interpretation and statistics.

To summarize, the long-lived royalactin-treated worms are more stress resistant and do not show any noticeable life-history trade-offs.

## Discussion

Many molecular biological studies of ageing focus on variants of a mechanism for which we here propose the descriptive term ‘defensive longevity’. In *C. elegans*, this includes reduced insulin/insulin-like growth factor signalling, reduced mitochondrial respiration and dietary restriction^15,44–46^. Due to the defensive mechanism, the proteome is stabilized and the generation of molecular damage minimized^8–10^. However, longevity attained as such is no clear guarantee for extended health^47–50^ and may bring along certain physiological trade-offs such as a smaller body volume^51^. Looking for possible alternative mechanisms that may escape this defensive strategy and may be characterized by more appealing health parameters, we studied the mode of action of royalactin, driven by the interesting queen bee phenotype associated with it^4^.

We found that royalactin treatment delays several ageing phenotypes (Fig. 1 & 5)^6,7^ and stimulates protein folding, mRNA translation and protein recycling (Fig. 1, 3 & 5) to promote longevity. Because this take on proteostasis in its very essence opposes the defensive mechanisms, we propose that royalactin induces an alternative, ‘copious’ approach to attain longevity: one involving proteome renewal via increased protein translation and degradation (Fig. 2 & 3). It is tempting to speculate that royalactin’s copious approach to proteostasis and longevity may be the reason for its seemingly improved stamina compared to other longevity models. However, further experiments are needed to substantiate mechanistic causality for healthspan, as we here show to be the case for longevity. Extensive information on the effect of royalactin in late-life *C. elegans* or in other organisms is lacking. It nevertheless seems that the increased health status may hold true for several readouts: we previously showed that royalactin-induced longevity depends on EGF(R) signalling^6^, and EGFR^20,52^ and EGF gain-of-function *C. elegans* also seem healthier (Monica Driscoll, personal communication). Our current findings should motivate increased scientific attention to the details of EGF(R) tuning in regard to healthy ageing.

A copious approach to longevity is interesting because it may prevent the accumulation of damage to proteins without necessarily reducing the incurrence of the damage itself. As the mono- and polysomes were elevated proportionally in royalactin-treated animals (Fig. 2), general translational capacity is increased, and we hypothesize that the differences observed in the proteome (Supplementary Dataset 1) stem from transcriptional changes, rather than from specificity at the translational level. This line of thinking resonates with the ‘turnover paradigm’ of earlier biogerontology literature [see ^53,54^ for brief overviews], but thus far, predominantly the opposite, defensive mechanism had been observed in long-lived animals^8,9^.

Now that experimental evidence for both is available, one may ask: which approach is the more advantageous one? In theory, each comes with its own drawbacks. When turnover is slowed down (defensive strategy), protein half-lives increase, in turn necessitating damage control over the existing proteome. Indeed, as prime examples of defensive longevity, *daf*-*2(*-) mutants activate various cytoprotective mechanisms, many of which are partially required for the lifespan-extending effect^55^. A benefit of lowering protein turnover is that the amount of erroneous proteins arising from protein synthesis is reduced and that cellular resources can thus be diverted away from *de novo* protein quality control towards somatic maintenance^10^. In the copious approach, when both protein synthesis and degradation are working fast, they need to be tightly coordinated as to avoid rapid proteome imbalance in case of desynchronization^56^. Upregulation of mRNA translation is expected, along with systems that help fold the nascent protein chains (*e.g*. co-translational chaperones) but also of systems enhancing breakdown of erroneous proteins resulting from *de novo* synthesis. Indeed, after royalactin treatment we observed a higher abundance of CCT chaperonins that help fold nascent protein chains in the cytosol (Supplementary Dataset 1), and a higher ability of the UPS to remove damaged/unstable proteins (Fig. 3). High protein turnover nevertheless remains an energy-costly process that may not be beneficial under all but the most beneficial environmental conditions. This may explain why copious longevity is likely much less prevalent in nature than defensive mechanisms are. Queen bees enjoy the luxury of being constantly fed a nutrient-rich mixture (including royalactin) by a horde of worker bees in a protected environment. As such, they are in a privileged position that may be permissive for the copious mechanism. When considering the lifestyle of several humans at risk of age-relate decline, it is not unreasonable to think this well-fed lifestyle more fitting than one generally associated with defensive longevity (e.g. dietary restriction). While the lifespan-extending effect of copious longevity in *C. elegans* is much more modest than that of defensive interventions, it is particularly appealing because of the observed health benefits.

Interventions other than royalactin may similarly induce copious longevity, which opens up interesting perspectives for future studies. For example, when *O. europaea*-derived products including the polyphenol tyrosol are added to the medium of *C. elegans* or of human fibroblasts, they induce proteomic and phenotypic effects similar to those of royalactin (Supplementary Table 1)^57–60^. This suggests a similar copious longevity mechanism, and we suspect more dietary and genetic interventions to follow suit in future discoveries.

In sum, we here provide evidence for a healthy ageing mechanism as induced by royalactin and depending on increased translation and proteasome function. As opposed to the better-known defensive mechanisms, we propose royalactin induces copious longevity. We could not yet detect any trade-offs with the increased lifespan, indicative of consistently positive effects on healthspan.

## Methods

### *C. elegans* strains

The following strains were obtained from the Caenorhabditis Genetic Center (CGC), which is funded by NIH Office of Research Infrastructure Programs (P40 OD010440): N2 WT Bristol, RB1206 *rsks*-*1(ok1255)*, UP233 *eor*-*1(cs28)*, AM141 *rmIs133[unc*-*54_p_::Q40::YFP]*, CL2070 *dvls70[hsp*-*16.2p::gfp;rol*-*6(su1006)]*, SJ4005 *zcls4[hsp*-*4p::gfp]*, SJ4100 *zcls13[hsp*-*6p::gfp]* and EU31 *skn*-*1(zu135)/nT1[unc*-*?(n754) let*-*?].* A four-times backcrossed RB1823 *gst*-*4(ok2358)* strain^61^ was kindly provided by Prof. Michael Miller. The strain PP608 (*hhIs64[unc*-*119(*+); *sur*-*5_p_::Ub^G76V^*-*GFP]*; *hhIs73[unc*-*119(*+); *sur*-*5_p_::mCherry])* was kindly provided by Prof. Thorsten Hoppe. We generated the LSC1208 *skn*-*1(zu135)/nT1[qIs51]* strain, by crossing EU31 animals with heterozygous *nT1(qIs51)* males. LSC1208 animals carry a chromosomal balancer with a pharyngeal GFP marker, making it easier to identify homozygous *skn*-*1(zu135)* animals via the absence of fluorescence rather than the absence of an *unc* phenotype (as in EU31 animals). Additionally, LSC1208 *skn*-*1(zu135)* homozygotes lay eggs that seldom hatch. The LSC1176 *eor*-*1(cs28)* strain was generated by outcrossing UP233 with WT, to a total of 5 times. Primers from a previous study^7^ were used to verify the *eor*-*1(cs28)* deletion allele via size estimation of PCR amplicons. Throughout the manuscript, the alleles *eor*-*1(cs28)*, *gst*-*4(ok2358)*, *rsks*-*1(ok1255)* and *skn*-*1(zu135)* are referred to as *eor*-*1(*-), *gst*-*4(*-) and *rsks*-*1(*-), respectively.

Worms were cultivated at 20°C on standard nematode growth medium (NGM) seeded with *E. coli* OP50, unless stated otherwise. NGM composition: 1.7% w/v agar, 50 mM NaCl, 0.50% w/v Bacto^™^ Peptone, 5 μg mL^-1^ cholesterol, 1 mM MgSO_4_, 1 mM CaCl_2_, 20 mM KH2PO4, 5 mM K_2_HPO_4_.

### Royalactin preparation

Royalactin (also known as Major Royal Jelly Protein 1 or MRJP1) was prepared from RJ via ultracentrifugation, according to ^6,7^ Royalactin-containing NGM plates were prepared by adding 1.7 μg mL^-1^ royalactin (unless stated otherwise) to liquid NGM immediately before pouring the plates.

### Differential proteomics: 2D-DIGE

WT N2 worms were synchronized by isolating eggs from gravid adults through hypochlorite treatment, as described before^6^. Synchronous L1 animals were placed on 9.5 cm diameter NGM or royalactin-containing (1.5 μg mL^-1^) NGM plates, both supplemented with 50 μg mL^-1^ streptomycin, and seeded with *E. coli* HB101. The HB101 strain is resistant to streptomycin. These conditions were used to ensure there was no variation in food source of the animals. Adding 120 μM of 5-fluoro-2’-deoxyuridine (FUdR) to the plates at the late L3 stage (circa 36 hours after plating out the L1 arrested larvae) avoided progeny production.

Sampling took place at day three of adulthood, six days after plating out the L1 larvae. Ten samples (biological replicates) per condition were collected, each consisting of approximately 2000 animals spread over six plates.

Animals were collected, washed and lysed and the resultant protein samples were differentially labelled and analysed using a two-dimensional difference gel electrophoresis (2D-DIGE) approach. See the Supplementary Methods for in-depth information.

### Network analysis

Gathered proteomics data was visualized using STRING (*<u>S</u>earch <u>T</u>ool for the <u>R</u>etrieval of <u>I</u>nteracting <u>G</u>enes/Proteins*, version 10.0), a database of known and predicted protein-protein interactions. All 35 unique protein identifications were used in each search. Settings for all networks: high confidence (>0.7) with clustering based on: *Co*-*expression*, *Experiments*, *Databases* and *Text mining.* See Supplementary Dataset 1B for GO and KEGG pathway analysis via STRING, and Supplementary Dataset 1C for tissue and phenotype enrichment analysis via WormBase^62^.

### *C. elegans* lifespan assays

Lifespan assays were performed similar to ^6^. In short, age-synchronized animals are transferred to NGM assay plates in the presence and absence of royalactin, and scored daily for survival. Animals are transferred to fresh assay plates every 3 to 4 days. Animals that do not move when gently prodded with a platinum wire are marked as dead, while animals that crawled off the plate or died due to vulval bursting or internal hatching were censored. The first day of adulthood, is always recorded as day 0. We used R^63^ to construct Kaplan-Meier survival curves, calculate mean and median lifespan and carry out all related statistical analyses. To compare survival between two conditions, a log-rank test was used (*i.e. survdiff* function of the *survival* package). The interaction of royalactin treatment between multiple conditions was compared using a Cox Proportional Hazard model (*i.e. coxph* function of the *survival* package). The corresponding p-values are referred to as *p_log-rank_* and *p_interaction_*. All resulting data can be found in Supplementary Dataset 2. For more detailed information regarding the setup and design of the lifespan assays, see the Supplementary Methods.

### Polysome profiling

In polysome profiling, differential centrifugation is used to separate the mRNA bound to multiple ribosomes and to single ribosomes (*i.e.* polysomes and monosomes) from translationally inactive mRNA^64,65^. By quantifying peak areas from the resulting RNA absorbance profiles, one can get an indication of the translational rate (*e.g*. fraction of translationally active mRNAs). Polysome profiling was performed similar to previous studies ^12,29^. Age synchronized WT animals were grown on NGM seeded with OP50 and collected at the first day of adulthood (55 hours post feeding L1 arrested larvae). Per sample, 200 μL of gently pelleted worms were lysed with an IKA T10 Ultra-Turrax homogenizer in 1 mL of lysis buffer (110 mM KAc, 10 mM MgAc_2_, 100 mM KCl, 10 mM MgCl_2_, 10 mM HEPES pH 7.6, 0.1% Tween-20, 2 mM DTT, 40 U/mL RNasin, 35.5 μM CHX). After 10 minutes on ice, worm extracts were centrifuged for 12 min at 16000 g and 4°C. The supernatant was transferred to Eppendorf LoBind DNA/RNA tubes, absorbance was measured with a Nanodrop spectrophotometer (Nanodrop ND-1000, Isogen Life Science) and samples were stored at −80°C.

Three separate centrifugation runs were performed, for a total of 14 samples. The day before a run, 10 – 55% sucrose gradients were prepared in gradient buffer (110 mM KAc, 10 mM MgAc_2_, 355 μM CHX, 10 mM HEPES pH 7.6) in 14 x 95 mm Ultra-Clear^™^ ultracentrifugation tubes (Beckman Coulter) and stored at 4°C. Each gradient was loaded with the same volume of worm extract (corresponding to 20 – 30 A_260nm_ units, Supplementary Dataset 3) followed by ultracentrifugation for a minimum of 90 min at 180000g and 4°C (Optima LE-80K Ultracentrifuge, Sw 40 Ti rotor, Beckman Coulter). As to not disturb the gradients, *slow acceleration* and *slow brake* were selected with a change to *no brake* for the final 1500 rpm. Gradients were collected via a Beckman Fraction Recovery system, by slowly displacing the gradients with 60% _w/v_ sucrose in gradient buffer using a peristaltic pump. The resulting droplets were collected in 96 well plates (UV-star, Greiner Bio-One, 4 droplets per well), until the entire sucrose gradient was transferred. Absorbance at 254 nm was measured using a Tecan Infinite M200 plate reader. Absorbance profiles were divided by the initial A_260nm_ units present in each sample (*i.e.* normalization for differential loading of total mRNA between samples) and are shown in Supplementary Dataset 3B. Gradients which were disturbed (*e.g*. due to the formation of air bubbles during collection) were discarded. Baseline-subtracted peak areas were calculated using GraphPad Prism 6, and statistically compared using a Student’s t-test with Benjamini-Hochberg multiple comparison correction ^66^ (reported as *p*_*t*-*test*, *FDR*_).

### *in vitro* proteasome activity

A 20S Proteasome Activity Assay kit (Merck, Chemicon APT280) was used to quantify hydrolysis of a fluorogenic proteasome substrate by *C. elegans* homogenates. Age synchronized L4 stage WT animals (48 hours post feeding L1 arrested larvae) were lysed with a IKA T10 Ultra-Turrax homogenizer in proteasome lysis buffer [50 mM HEPES, 150 mM NaCl, 5 mM EDTA, 2 mM ATP, 1 mM DTT, pH 7.5; similar to refs.^38^,^67^]. After centrifugation (18000 g, 15 min, 4°C), supernatants were transferred to Protein LoBind tubes, on ice. Protein concentrations were measured using a Qubit assay (Invitrogen) with Bovine Serum Albumin protein standards prepared in proteasome lysis buffer.

Per sample, 7.5 μg of protein in 20 μL proteasome lysis buffer was mixed with 500 μM of the chymotryptic substrate Suc-LLVY-AMC (in DMSO), proteasome assay buffer (25 mM HEPES, 5 mM EDTA, 0.5% NP-40, 0.01% SDS _w/v_, pH 7.5; similar to ref.^38^) and DMSO or proteasome inhibitor (7.1 μM lactacystin in DMSO). The assay was performed according to the manufacturer’s instructions, in a 96 well plate with randomized layout, using two technical replicates per sample and 8 biological replicates per condition. For half of the experimental conditions, lactacystin was added. Fluorescence was measured at 37°C during 90 min (Flexstation II, Molecular Devices, λ_ex_/λ_em_ = 380/460 nm). Background signals due to 7-Amino-4-methylcoumarin (AMC) auto hydrolysis were subtracted. An AMC standard curve was constructed to convert relative fluorescent units (RFU) into a substrate hydrolysis rate estimate (μM h^-1^).

### *in vivo* 20S proteasome activity

To quantify proteasomal activity *in vivo* we used the PP608 strain which broadly expresses GFP fused to a mutant form of ubiquitin (Ub^G76V^-GFP) via the near-ubiquitous *sur*-*5* promoter. This irreversibly targets GFP for proteasomal proteolysis^38,68,69^. The decrease in GFP fluorescence intensity reflects the global Ub^G76V^-GFP turnover rate and correlates directly with the rate of proteasomal degradation. The strain also contains a more stable red fluorescent reporter (mCherry) driven by the same broadly expressed promoter^69^. By taking both the GFP and mCherry fluorescent intensities into account, one can rule out differences in GFP signal due to differences in transgene expression and protein translation. Low Ub^G76V^-GFP/mCherry levels reflect enhanced breakdown of the ubiquitin-tagged GFP by the proteasome, therefore a higher ability of the UPS to remove damaged and unstable proteins^20,38^.

The N-terminally linked ubiquitin moiety contains a G76V substitution which prevents cleavage by ubiquitin hydrolases, leading to polyubiquitination of GFP by E3 ligases^20,70^. In essence, GFP is irreversibly targeted for proteasomal proteolysis. Fluorescence was quantified analogous to the automated method described in Detienne et al.^7^, using a microplate reader to measure thousands of nematodes simultaneously (Flexstation II, Molecular Devices). Background-corrected fluorescence intensities were calculated for both the Ub^G76V^-GFP (λ_ex_/λ_em_ = 470/509 nm) and mCherry signals (λ_ex_/λ_em_ = 590/630 nm) at multiple time points. Animals were grown on regular or royalactin-containing NGM supplemented with 115 μM FUdR from the L4 stage onwards, until the time of measurement. Measurements were conducted on multiple independent populations, pooled and statistically compared using a Student’s t-test with Benjamini-Hochberg multiple comparison correction^66^.

### Reproductive fitness

To measure reproductive fitness, the protocol of Keith et al.^71^ was followed with minor modifications. For each condition, 2500 L1 arrested WT animals (age-synchronized via hypochlorite treatment) were transferred to 9 cm diameter NGM plates, in the presence and absence of royalactin (defined as t = 0). After reaching the L4 stage, hermaphrodites were transferred individually to 3.5 cm diameter plates and allowed to self-fertilize. Per condition, 24 plates were used with one animal (biological replicate) on each plate. During the first three days, animals were transferred thrice a day to fresh plates (at 9h, 15h and 21h) taking care not to transfer any eggs. Nearing the end of the experiment, the interval between transfers was increased: adult animals were only transferred twice or once daily until the end of the egg-laying period. The amount of hatched progeny was manually counted on all the plates, using a binocular microscope. The progeny of two parent animals that died or went missing during the first ten time points (*i.e.* within 126 hours post L1 feeding; one animal per condition) were excluded from the analysis. A repeated measures two-way ANOVA with Sidak’s correction for multiple comparisons was used to compare the amount of progeny at multiple time points (using the raw data), and a Mann-Whitney test to compare the total amount of progeny. The reproductive rate (amount of progeny day^-1^) was calculated analogous to Muschiol et al.^72^, and is shown in addition to the raw data (Supplementary Dataset 4).

### Body size measurements

Body size of WT adults was quantified analogous to refs.^73^ and ^74^. Synchronized L1 worms were grown on NGM or royalactin-containing NGM plates. After 48 hours, FUdR was added at a final concentration of 135 μM. After an additional 26 hours, the worm populations were collected and washed with S Basal (0.1 M NaCl, 44 mM KH_2_PO_4_, 6 mM K_2_HPO_4_) and fixed in 4% _w/v_ formaldehyde. The average length (L) and width (W) of individual worms was analyzed with a RapidVUE Particle Shape and Size Analyzer (Beckman Coulter). Each of the six biological replicates (3 per condition) was run at least three times (technical replicates). Worm volumes were approximated using the cylinder volume formula: V = πL(W/2)^2^.

Averages were statistically compared using a Student’s t-test with Benjamini-Hochberg multiple comparison correction^66^.

### Reductive stress (DTT) assays

Royalactin-treated and untreated adult worms were subjected to reductive stress via incubation in 5 mM dithiothreitol (DTT,) as described by Glover-Cutter et al.^75^. Synchronous populations were obtained by timed egg-lay and treated with 135 μM FUdR at the young adult stage. At the start of each assay (i.e. at day 1 or 3 of adulthood), worms were collected and washed with M9 buffer (85 mM NaCl, 22 mM KH_2_PO_4_, 41 mM Na_2_HPO_4_, 17 mM NH4Cl) followed by 24h incubation in 1.7 mL of 5 mM DTT in M9 buffer in Eppendorf tubes while rotating. After incubation, the stressed worms were plated out on NGM plates containing a lawn of OP50 *E. coli* bacteria and immediately scored for survival.

### Heat stress assays

Royalactin-treated and untreated animals were subjected to heat stress, in essence as described in Zevian et al.^76^. Synchronous WT populations were obtained by hypochlorite treatment and grown on NGM at 20°C before switching to a pre-heated 35°C incubator (at either the young adult stage or at day 6 of adulthood). Worms were briefly taken out every hour to assay their survival, until no more animals remained. Worms were scored and analyzed as described for lifespan assays. For the animals assayed at day 6 of adulthood, FUdR was added to the plates to maintain a synchronous population. Our observation that these older adults are more tolerant to heat stress (Fig. 5B), is in line with observations in previous studies^76,77^.

### Proteotoxic aggregation assays

We examined polyQ aggregation in transgenic AM141 animals^78^ at multiple time points. At 48 hours post L1 feeding, FUdR was added to the culture plates for both conditions (*i.e.* royalactin-treated and untreated). We used a Zeiss Axio Imager Z1 microscope (10x objective) to acquire images similar to ^7^, and ImageJ v1.48^79^ to outline and quantify the amount of fluorescent aggregates of individual worms (using the *Find Maxima* function with a noise tolerance of 10). Each replicate consists of an individual worm. The laser strength and exposure time were held constant for all data acquisition.

### Oxidative stress (paraquat) assays

Paraquat-induced oxidative stress assays were performed similar to ^80–82^. L4 stage worms were transferred from NGM plates to NGM plates containing 8 mM paraquat (Acros Organics, Belgium). All plates were seeded with *E. coli* HT115(DE3) and contained 50 μg mL^-1^ ampicillin to avoid bacterial contamination, which otherwise could skew results (*e.g*. enhanced sensitivity of some mutants to bacterial contamination). Survival was checked daily and analyzed as described for lifespan assays, without the use of FUdR. *eor*-*1(*-) refers to the LSC1176 strain, *gst*-*4(*-) to the RB1823 strain and *skn*-*1(*-) to *skn*-*1(zu135)* homozygotes of the LSC1208 strain.

### Transcriptional reporter strains

GFP fluorescence of the *C. elegans* transcriptional reporter strains, CL2070 *(hsp*-*16.2p::gfp)*, SJ4005 (*hsp*-*4p::gfp*) and SJ4100 (*hsp*-*6p::gfp*) was quantified using an automated method, as described in Detienne et al.^7^. Animals were grown without FUdR in the presence and absence of royalactin on NGM plates seeded with OP50, until the late L4 stage, before transfer to a 96 well plate for fluorescence quantification. Pooled data from multiple experiments is shown.

## Acknowledgements

GD and PVdW are IWT Flanders research fellows, WDH is an FWO Flanders postdoctoral researcher and BC benefits from an FWO SB mandate. The authors wish to thank KU Leuven (C14/45/049), the European Research Council (H2020 grant 633589) and FWO Flanders (G043710N, G069713, G095915, G052217N) for financial support. Some strains were provided by the CGC, which is funded by NIH Office of Research Infrastructure Programs (P40 OD010440). We are grateful to Prof. Michael Miller and Prof. Thorsten Hoppe for sharing additional strains, and to Luc Vanden Bosch for providing excellent technical support. We would like to acknowledge the following persons for providing valuable feedback on experimental design and methods: Kira Glover-Cutter (Blackwell lab), Joshi Kishore (Rongo lab), Sarah Tremmel (Antebi lab) and Noa Liberman-Isakov (Sinclair/Greer lab).

## Author contributions

Conceived and designed the experiments: GD assisted by WDH, BPB, LS and LT. Performed the experiments: GD assisted by PVdW and BC. Analyzed the data: GD assisted by PVdW and WDH. Interpreted the data: GD assisted by all co-authors. Wrote the manuscript: GD, WDH and LT, assisted by PVdW, PBP and LS.

## References

1. de Magalhães, J. P., Costa, J. & Church, G. M. An analysis of the relationship between metabolism, developmental schedules, and longevity using phylogenetic independent contrasts. J Gerontol A Biol Sci Med Sci 62, 149–160 (2007).

2. Hansen, M., Flatt, T. & Aguilaniu, H. Reproduction, fat metabolism, and life span: What is the connection? Cell Metab. 17, 10–19 (2013).

3. Page, R. E. & Peng, C. Y. Aging and development in social insects with emphasis on the honey bee, Apis mellifera L. Exp. Gerontol. 36, 695–711 (2001).

4. Kamakura, M. Royalactin induces queen differentiation in honeybees. Nature 473, 478–483 (2011).

5. Fan, P. et al. Functional and proteomic investigations reveal Major Royal Jelly Protein 1 associated with anti-hypertension activity in mouse vascular smooth muscle cells. Sci. Rep. 6, 30230 (2016).

6. Detienne, G., De Haes, W., Ernst, U. R., Schoofs, L. & Temmerman, L. Royalactin extends lifespan of *Caenorhabditis elegans* through epidermal growth factor signaling. Exp. Gerontol. 60, 129–135 (2014).

7. Detienne, G., Van de Walle, P., De Haes, W., Schoofs, L. & Temmerman, L. SKN-1-independent transcriptional activation of glutathione S-transferase 4 (GST-4) by EGF signaling. Worm 5, e1230585 (2016).

8. Depuydt, G., Shanmugam, N., Rasulova, M., Dhondt, I. & Braeckman, B. P. Increased protein stability and decreased protein turnover in the *Caenorhabditis elegans* Ins/IGF-1 daf-2 mutant. Journals Gerontol. - Ser. A Biol. Sci. Med. Sci. 71, 1553–1559 (2016).

9. Dhondt, I. et al. FOXO/DAF-16 Activation Slows Down Turnover of the Majority of Proteins in *C. elegans*. Cell Rep. 16, 3028–3040 (2016).

10. Vellai, T. & Takács-Vellai, K. Regulation of protein turnover by longevity pathways. Adv. Exp. Med. Biol. 694, 69–80 (2010).

11. Syntichaki, P., Troulinaki, K. & Tavernarakis, N. eIF4E function in somatic cells modulates ageing in *Caenorhabditis elegans*. Nature 445, 922–926 (2007).

12. Pan, K. Z. et al. Inhibition of mRNA translation extends lifespan in *Caenorhabditis elegans*. Aging Cell 6, 111–119 (2007).

13. Henderson, S. T., Bonafè, M. & Johnson, T. E. daf-16 protects the nematode *Caenorhabditis elegans* during food deprivation. J. Gerontol. A. Biol. Sci. Med. Sci. 61, 444–60 (2006).

14. Curran, S. P. & Ruvkun, G. Lifespan regulation by evolutionarily conserved genes essential for viability. PLoS Genet. 3, e56 (2007).

15. Hansen, M. et al. Lifespan extension by conditions that inhibit translation in *Caenorhabditis elegans*. Aging Cell 6, 95–110 (2007).

16. Tavernarakis, N. Ageing and the regulation of protein synthesis: a balancing act? Trends Cell Biol. 18, 228–235 (2008).

17. Smith, E. D. et al. Quantitative evidence for conserved longevity pathways between divergent eukaryotic species. Genome Res. 18, 564–570 (2008).

18. Essers, P. et al. Reduced insulin/insulin-like growth factor signaling decreases translation in Drosophila and mice. Sci. Rep. 6, 30290 (2016).

19. Kaeberlein, M. & Kennedy, B. K. Hot topics in aging research: protein translation and TOR signaling, 2010. Aging Cell 10, 185–90 (2011).

20. Liu, G., Rogers, J., Murphy, C. T. & Rongo, C. EGF signalling activates the ubiquitin proteasome system to modulate *C. elegans* lifespan. EMBO J. 30, 2990–3003 (2011).

21. Mendoza, M. C., Er, E. E. & Blenis, J. The Ras-ERK and PI3K-mTOR pathways: cross-talk and compensation. Trends Biochem. Sci. 36, 320–328 (2011).

22. McQuary, P. R. et al. *C. elegans* S6K Mutants Require a Creatine-Kinase-like Effector for Lifespan Extension. Cell Rep. 14, 2059–2067 (2016).

23. Dong, M.-Q. et al. Quantitative mass spectrometry identifies insulin signaling targets in *C. elegans*. Science (80-.). 317, 660–3 (2007).

24. Li, X. et al. Specific SKN-1/NrF stress responses to perturbations in translation elongation and proteasome activity. PLoS Genet. 7, 9–11 (2011).

25. Reis-Rodrigues, P. et al. Proteomic analysis of age-dependent changes in protein solubility identifies genes that modulate lifespan. Aging Cell 11, 120–7 (2012).

26. Ennis, H. L. & Lubin, M. Cycloheximide: Aspects of Inhibition of Protein Synthesis in Mammalian Cells. Science (80-.). 146, 1474–1476 (1964).

27. Schneider-Poetsch, T. et al. Inhibition of eukaryotic translation elongation by cycloheximide and lactimidomycin. Nat. Chem. Biol. 6, 209–217 (2010).

28. Takauji, Y. et al. Restriction of protein synthesis abolishes senescence features at cellular and organismal levels. Sci. Rep. 6, 18722 (2016).

29. Stout, G. J. et al. Insulin/IGF-1-mediated longevity is marked by reduced protein metabolism. Mol. Syst. Biol. 9, 679–679 (2014).

30. Heyer, E. E. & Moore, M. J. Redefining the Translational Status of 80S Monosomes. Cell 164, 757–769 (2016).

31. López-Otín, C., Blasco, M., Partridge, L., Serrano, M. & Kroemer, G. The hallmarks of aging. Cell 153, 1194–217 (2013).

32. Schmidt, M. & Finley, D. Regulation of proteasome activity in health and disease. Biochim. Biophys. Acta 1843, 13–25 (2014).

33. Chondrogianni, N., Georgila, K., Kourtis, N., Tavernarakis, N. & Gonos, E. S. 20S proteasome activation promotes life span extension and resistance to proteotoxicity in *Caenorhabditis elegans*. FASEB J. 29, 611–622 (2015).

34. Chondrogianni, N., Sakellari, M., Lefaki, M., Papaevgeniou, N. & Gonos, E. S. Proteasome activation delays aging in vitro and in vivo. Free Radic. Biol. Med. 71, 303–320 (2014).

35. Pride, H. et al. Long-lived species have improved proteostasis compared to phylogenetically-related shorter-lived species. Biochem. Biophys. Res. Commun. 457, 669–675 (2015).

36. Ellina, M., Bouris, P., Kletsas, D., Aletras, A. J. & Karamanos, N. K. EGF/EGFR signaling axis is a significant regulator of the proteasome expression and activity in colon cancer cells. Sci. Res. 1–10 (2014). doi:10.14293/A2199-1006.01.SOR-LIFE.AC0E6.v1

37. Bogyo, M. & Wang, E. W. Proteasome inhibitors: complex tools for a complex enzyme. Curr. Top. Microbiol. Immunol. 268, 185–208 (2002).

38. Joshi, K. K., Matlack, T. L. & Rongo, C. Dopamine signaling promotes the xenobiotic stress response and protein homeostasis. EMBO J. 35, 1–17 (2016).

39. Luo, S., Shaw, W. M., Ashraf, J. & Murphy, C. T. TGF-ß Sma/Mab Signaling Mutations Uncouple Reproductive Aging from Somatic Aging. PLoS Genet. 5, e1000789 (2009).

40. Hughes, S. E., Evason, K., Xiong, C. & Kornfeld, K. Genetic and pharmacological factors that influence reproductive aging in nematodes. PLoS Genet. 3, 0254–0265 (2007).

41. Shen, P. et al. Piceatannol extends the lifespan of *Caenorhabditis elegans* via DAF-16. Biofactors 43, 379–387 (2017).

42. Dues, D. J. et al. Aging causes decreased resistance to multiple stresses and a failure to activate specific stress response pathways. Aging (Albany. NY). 8, 777–95 (2016).

43. Fan, H.-C. et al. Polyglutamine (PolyQ) diseases: genetics to treatments. Cell Transplant. 23, 441–58 (2014).

44. Barzilai, N., Crandall, J. P., Kritchevsky, S. B. & Espeland, M. A. Metformin as a Tool to Target Aging. Cell Metab. 23, 1060–1065 (2016).

45. Baker, B. M., Nargund, A. M., Sun, T. & Haynes, C. M. Protective Coupling of Mitochondrial Function and Protein Synthesis via the eIF2α Kinase GCN-2. PLoS Genet. 8, e1002760 (2012).

46. Depuydt, G. et al. Reduced insulin/insulin-like growth factor-1 signaling and dietary restriction inhibit translation but preserve muscle mass in *Caenorhabditis elegans*. Mol. Cell. Proteomics 12, 3624–39 (2013).

47. Hansen, M. & Kennedy, B. K. Does Longer Lifespan Mean Longer Healthspan? Trends Cell Biol. 26, 565–568 (2016).

48. Podshivalova, K., Kerr, R. A. & Kenyon, C. How a Mutation that Slows Aging Can Also Disproportionately Extend End-of-Life Decrepitude. Cell Rep. 19, 441–450 (2017).

49. Ewald, C. Y., Castillo-Quan, J. I. & Blackwell, T. K. Untangling Longevity, Dauer, and Healthspan in *Caenorhabditis elegans* Insulin/IGF-1-Signalling. Gerontology 64, 96–104 (2018).

50. Bansal, A., Zhu, L. J., Yen, K. & Tissenbaum, H. a. Uncoupling lifespan and healthspan in *Caenorhabditis elegans* longevity mutants. Proc. Natl. Acad. Sci. 112, E277–E286 (2015).

51. Iser, W. B. & Wolkow, C. A. DAF-2/insulin-like signaling in *C. elegans* modifies effects of dietary restriction and nutrient stress on aging, stress and growth. PLoS One 2, e1240 (2007).

52. Iwasa, H., Yu, S., Xue, J. & Driscoll, M. Novel EGF pathway regulators modulate *C. elegans* healthspan and lifespan via EGF receptor, PLC-gamma, and IP3R activation. Aging Cell 9, 490–505 (2010).

53. Braeckman, B. P. & Vanfleteren, J. R. Genetic control of longevity in *C. elegans*. Exp. Gerontol. 42, 90–8 (2007).

54. Ryazanov, A. G. & Nefsky, B. S. Protein turnover plays a key role in aging. Mech. Ageing Dev. 123, 207–13 (2002).

55. Shore, D. E., Carr, C. E. & Ruvkun, G. Induction of Cytoprotective Pathways Is Central to the Extension of Lifespan Conferred by Multiple Longevity Pathways. PLoS Genet. 8, e1002792 (2012).

56. Visscher, M. et al. Proteome-wide Changes in Protein Turnover Rates in *C. elegans* Models of Longevity and Age-Related Disease. Cell Rep. 16, 3041–3051 (2016).

57. Cañuelo, A. et al. Tyrosol, a main phenol present in extra virgin olive oil, increases lifespan and stress resistance in *Caenorhabditis elegans*. Mech. Ageing Dev. 133, 563–574 (2012).

58. Cañuelo, A. & Peragón, J. Proteomics analysis in *Caenorhabditis elegans* to elucidate the response induced by tyrosol, an olive phenol that stimulates longevity and stress resistance. Proteomics 13, 3064–3075 (2013).

59. Haris Omar, S. Oleuropein in Olive and its Pharmacological Effects. Sci. Pharm. 78, 133–154 (2010).

60. Katsiki, M., Chondrogianni, N., Chinou, I., Rivett, A. J. & Gonos, E. S. The olive constituent oleuropein exhibits proteasome stimulatory properties in vitro and confers life span extension of human embryonic fibroblasts. Rejuvenation Res. 10, 157–72 (2007).

61. Edmonds, J. W. et al. Insulin/FOXO Signaling Regulates Ovarian Prostaglandins Critical for Reproduction. Dev. Cell 19, 858–871 (2010).

62. Angeles-Albores, D., N Lee, R. Y., Chan, J. & Sternberg, P. W. Tissue enrichment analysis for C. elegans genomics. BMC Bioinformatics 17, 366 (2016).

63. R Development Core Team. R: A language and environment for statistical computing. R Foundation for Statistical Computing, Vienna, Austria. ISBN 3–900051-07–0, URL http://www.R-project.org/. (2013).

64. Chassé, H., Boulben, S., Costache, V., Cormier, P. & Morales, J. Analysis of translation using polysome profiling. Nucleic Acids Res. gkw907 (2016). doi:10.1093/nar/gkw907

65. Mašek, T., Valášek, L. & Pospíšek, M. Polysome analysis and RNA purification from sucrose gradients. Methods Mol. Biol. 703, 293–309 (2011).

66. Benjamini, Y. & Hochberg, Y. Controlling the false discovery rate: a practical and powerful approach to multiple testing. J. R. Stat. Soc. Ser. B 57, 289–300 (1995).

67. Meng, F., Li, J., Wang, W. & Fu, Y. Gengnianchun, a Traditional Chinese Medicine, Enhances Oxidative Stress Resistance and Lifespan in *Caenorhabditis elegans* by Modulating daf-16/FOXO. Evidence-*Based Complement. Altern. Med.* 2017, 1–10 (2017).

68. Keith, S. A. et al. Graded Proteasome Dysfunction in Caenorhabditis elegans Activates an Adaptive Response Involving the Conserved SKN-1 and ELT-2 Transcription Factors and the Autophagy-Lysosome Pathway. PLoS Genet. 12, 1–39 (2016).

69. Segref, A., Torres, S. & Hoppe, T. A screenable in vivo assay to study proteostasis networks in *Caenorhabditis elegans*. Genetics 187, 1235–1240 (2011).

70. Stack, J. H., Whitney, M., Rodems, S. M. & Pollok, B. a. A ubiquitin-based tagging system for controlled modulation of protein stability. Nat. Biotechnol. 18, 1298–1302 (2000).

71. Keith, S. A., Amrit, F. R. G., Ratnappan, R. & Ghazi, A. The *C. elegans* healthspan and stress-resistance assay toolkit. Methods 68, 476–486 (2014).

72. Muschiol, D., Schroeder, F. & Traunspurger, W. Life cycle and population growth rate of *Caenorhabditis elegans* studied by a new method. BMC Ecol. 9, 14 (2009).

73. Van Sinay, E. et al. Evolutionarily conserved TRH neuropeptide pathway regulates growth in *Caenorhabditis elegans*. Proc. Natl. Acad. Sci. 114, E4065–E4074 (2017).

74. De Haes, W. et al. Metformin promotes lifespan through mitohormesis via the peroxiredoxin PRDX-2. Proc. Natl. Acad. Sci. 1–9 (2014). doi:10.1073/pnas.1321776111

75. Glover-Cutter, K. M., Lin, S. & Blackwell, T. K. Integration of the Unfolded Protein and Oxidative Stress Responses through SKN-1/Nrf. PLoS Genet. 9, (2013).

76. Zevian, S. C. & Yanowitz, J. L. Methodological considerations for heat shock of the nematode *Caenorhabditis elegans*. Methods 68, 450–7 (2014).

77. Angeli, S. et al. A DNA synthesis inhibitor is protective against proteotoxic stressors via modulation of fertility pathways in *Caenorhabditis elegans*. Aging (Albany. NY). 5, 759–769 (2013).

78. Morley, J. F., Brignull, H. R., Weyers, J. J. & Morimoto, R. I. The threshold for polyglutamine-expansion protein aggregation and cellular toxicity is dynamic and influenced by aging in *Caenorhabditis elegans*. Proc. Natl. Acad. Sci. U. S. A. 99, 10417–22 (2002).

79. Schneider, C. A., Rasband, W. S. & Eliceiri, K. W. NIH Image to ImageJ: 25 years of image analysis. Nat. Methods 9, 671–675 (2012).

80. Zarse, K. et al. Impaired Insulin/IGF1 Signaling Extends Life Span by Promoting Mitochondrial L-Proline Catabolism to Induce a Transient ROS Signal. Cell Metab. 15, 451–465 (2012).

81. Zheng, S. et al. Mulberry leaf polyphenols delay aging and regulate fat metabolism via the germline signaling pathway in *Caenorhabditis elegans*. Age (Omaha). 36, 9719 (2014).

82. Wilson, M. A. et al. Blueberry polyphenols increase lifespan and thermotolerance in *Caenorhabditis elegans*. Aging Cell 5, 59–68 (2006).

